# A missing link in mutualistic networks: symbiotic fungi in plant-animal interactions

**DOI:** 10.1101/761270

**Authors:** Priscila Chaverri, Gloriana Chaverri

**Author notes:** Name and complete mailing address of the person to whom correspondence should be sent*: Priscila Chaverri, Escuela de Biología, Universidad de Costa Rica, San Pedro, Costa Rica. Statement of authorship*: PC and GC conceived the study, PC and GC designed the study, PC worked with the fungi, GC worked with the bats, PC and GC analyzed data; PC and GC wrote the paper. Data accessibility*: Data is available upon request: Raw and filtered data from metabarcoding, including results from OTU clustering, taxonomic classification, abundance tables and ITS sequences from each OTU.

## Abstract

We explored the hypothesis of an indirect mutualistic relationship (i.e., when the association between two species is modified by a third one) within a plant-animal seed dispersal network. Bats are important long-distance dispersers of many tropical plants, yet, by consuming fruits they may disperse not only the plant’s seeds, but also the endosymbiotic fungi within those fruits. We characterized fungal communities in fruits of *Ficus colubrinae* and in feces of *Ectophylla alba* to determine if passage through the digestive tract of the bats affected the total mycobiome. Results show a significant reduction, after passage through the gut, of fungi known to be plant pathogenic, while abundance of species known to have beneficial properties significantly increased. These findings suggest that the role of frugivores in plant-animal mutualistic networks may extend beyond seed dispersal: they also promote the dispersal of potentially beneficial microbial symbionts while hindering those that can cause plant disease.

## INTRODUCTION

Ecological networks have been the topic of extensive research, yet it seems that we still have important missing pieces of these complex puzzles (Olesen *et al.* 2011). For example, few studies have addressed the role of parasites in networks (Lafferty *et al.* 2008), and indirect interactions —those that occur when the interaction between two species is modified by a third one (Levine 1976; Holt 1977)— are seldom addressed and are still poorly understood (Miller & TerHorst 2012; Michalet *et al.* 2015). Notwithstanding, indirect interactions are ubiquitous in ecological networks and play a critical role in shaping the bonds among species within communities (Strauss 1991; Wootton 1994; Irwin 2006; Blanc & Walters 2008; Poelman *et al.* 2011). While we typically believe that the survival of a species depends solely on the protection of its direct interactions, realistically it may also depend on how those interactions are shaped by species indirectly linked to each node in the network; the loss of any of these indirect links may have unforeseen effects on network functioning. Therefore, in light of the vulnerability of many species due to human activities, such as introduction of exotic species, habitat loss, introduction of parasites, and climate change, we need to acknowledge both the direct and indirect interactions of a taxon within ecological networks to fully understand the potential effects of its loss.

A type of ecological network that has received much attention is the one that encompasses interactions between plants and their animal pollinators and seed dispersers; the interaction is mutually beneficial because animals help transport pollen and seeds, and in exchange, obtain food (reviewed in (Bascompte 2009). Studies of plant-animal mutualistic networks have focused on understanding the relationship among interacting species (Petanidou *et al.* 2008; Carnicer *et al.* 2009), the importance of particular taxa and network topology for maintaining network resilience (Okuyama & Holland 2008; Bascompte & Stouffer 2009; Mello *et al.* 2011; Allesina & Tang 2012), and how the topology of mutualistic networks may influence resilience (Okuyama & Holland 2008; Bascompte & Stouffer 2009; Allesina & Tang 2012), and even species diversity (Bascompte & Jordano 2007; Allesina & Tang 2012). However, these studies have neglected an important component of plant communities: their microbial epi- and endosymbionts (endo = inside, epi = on the surface; symbiosis = two different organisms living in close physical association).

A group of important microbial symbionts are endophytic fungi, which live within aerial tissues of plants without causing any visible negative impact. Even though some endophytes may be latent pathogens or saprotrophs (Porras-Alfaro & Bayman 2011), in many cases these endosymbionts provide benefits to the plant, including protection against diseases and pests, plant growth, and reduction of drought stress (Rodriguez *et al.* 2009). As a result, endophytic fungi are regarded as critical components of any healthy plant community. Less is known about epiphytic or phyllosphere fungi, but there is also evidence of benefits to the host plant (Andrews 1992; Lindow 2006).

The ability of a fungal species to disperse its spores (or other propagules such as hyphae, chlamydospores, and fruiting structures) is one of the factors that influence fungal diversity in a natural ecosystem (Kohn 2005; Persoh 2015). Fungi can only disperse their spores short distances (a few centimeters at the most) using their own means (e.g., forcible ejection or discharge) (Roper *et al.* 2008; Galante *et al.* 2011). Consequently, they rely on other ways for long distance dispersal, e.g., water, wind, and animals. Wind has been reported as the most efficient way to disperse fungal spores long distances. However, in natural forests, especially old-growth, wind may not have a large influence in spore dispersal because trees and understory vegetation provide a barrier to wind movement (Chen *et al.* 1993; Milleron *et al.* 2012). Therefore, it is expected that other factors besides wind are influencing long-distance dispersal of fungi in natural tropical forests. Considering that animals are capable of long-distance dispersal of plant seeds, it is possible that their role extends to the dispersal of fungal spores and other propagules.

In this study, we aim to explore the hypothesis of an indirect mutualistic relationship between fruit-eating animals, specifically bats, and symbiotic fungi (with emphasis on endophytes) that grow within the tissues of fruits that bats eat. Many animals may disperse fungi directly by eating mushrooms and then defecating the spores; or indirectly, by eating other plant parts that contain these fungi. The direct consumption of fungal fruiting bodies, or mycophagy, and spore dispersal has been described several times in insects, rodents, marmosets, and other mammals (Johnson 1996; Epps & Arnold 2010; Hilario & Ferrari 2010). In some cases it was reported that fungal spores survive and their germination is improved after passing the digestive tract of truffle-eating rodents or other ground-dwelling animals (Johnson 1996). However, indirect fungal propagule dispersal is woefully unknown. We use bats as a model to understand this interaction because these mammals are important long-distance dispersers of many tropical plants (Kunz *et al.* 2011). Yet by consuming fruits, bats may disperse not only the plant’s seeds, but also the fungi that are contained in those fruits. Bats may be particularly good dispersers of fungi because they fly long distances each night, defecate during flight, may retain viable propagules for long periods of time, and because unlike birds, fruit-eating bats often venture into deforested areas that may otherwise lack input of beneficial fungal spores (Cardoso DaSilva *et al.* 1996; Medellin & Gaona 1999; Shilton *et al.* 1999; Dumont 2003). This study represents a step towards identifying an interaction that may have consequences for the preservation of healthy tropical ecosystems, because endophytic fungi provide important benefits for host plants, improving plant fitness and thus increasing vital resources for the entire fruit-eating community.

Since fungi can develop in any plant tissue as endophytes, including fruits (Martinson *et al.* 2012; Tiscornia *et al.* 2012), it is presumed that bats may disperse fungal propagules that are consumed from these structures. To explore this completely unknown hypothesis that bats are also long-distance dispersers of endophytic fungi, many questions need to be answered. First, we need to determine which species of endophytic fungi are present in fruit tissues, and whether the same species are present in bat feces. In this project, we aim to answer these basic questions, yet many interrogations will remain regarding the relationship between frugivores, plants, and endophytic fungi. We hope our answers will begin to shed light on this potential interaction and hopefully foster further scrutiny.

## MATERIALS AND METHODS

### Study system and site

Fresh ripe fig fruits (*Ficus colubrinae* Standl., Moraceae) and Honduran White Bat (*Ectophylla alba* Allen, Chiroptera: Phyllostomidae) fecal samples were collected for fungal community analyses. It has been reported that *E. alba* feeds almost exclusively from *F. colubrinae* fruits (Brooke 1990). *Ectophylla alba* is known only from Honduras, Nicaragua, Costa Rica and western Panama (Reid 1997). In 2008, IUCN elevated this bat species to a near threatened Red List category (Rodríguez-Herrera & Pineda 2015). Populations of this bat species have been declining due to urbanization and strong habitat (e.g., *Heliconia* leaves from intermediate secondary succession forests for roosting) and diet preferences (i.e., *F. colubrinae* fruits) (Rodríguez-Herrera *et al.* 2008; Rodríguez-Herrera & Pineda 2015). *Ficus colubrinae* is an understory tree distributed from Mexico to Colombia and fruits throughout the year (La Selva Florula Digital, http://sura.ots.ac.cr/florula4/). Fruit and fecal samples were collected in La Selva Biological Station (Sarapiquí, Heredia, Costa Rica).

### Fruit and fecal sample collection

We attempted to capture bats and collect fruits from the same trees, as to increase the chances that the fecal samples came from the fruits consumed in that tree. Two trees that were fruiting at the time of the fieldwork (February 2015) were chosen for fruit samples, to place mist-nets and capture bats. Fruits were collected and placed in small Ziploc bags. Ten ripe fruits were collected for total mycobiome (endo- and epiphytic fungi) analyses (targeted-amplicon metagenomics or metabarcoding). The fruits for metabarcoding analyses were placed in 2-mL Eppendorf microtubes with silica gel for posterior DNA extraction.

Thirteen bats were captured with mist nets (Ecotone, Poland) and immediately placed in sterilized cloth bags. To avoid cross-contamination, we cleaned our hands with an alcohol-based hand gel before releasing every bat. When bats defecated, sterilized cotton swabs were used to obtain the fecal sample from the cloth bag. The approximate volume collected was 1/2 “pea-size” amount. Fecal samples were placed in 2-mL Eppendorf tubes, in Ziploc bags with silica gel, and then placed in the freezer for subsequent metabarcoding analyses.

### Molecular work

Genomic DNA from fruits was extracted with the following protocol: fruits were placed in the −20 °C freezer for at least one day and then, when ready to extract, placed into new tubes prefilled with 500 μm garnet beads and a 6-mm zirconium grinding satellite bead (OPS Diagnostics LLC, NJ, U.S.A.). For fruit tissue homogenization, a FastPrep® instrument (Zymo Research, Irvine, CA, U.S.A.) was used. 750 μl of Qiagen® Lysis Buffer AP1 and 6 ul of QiaGen® RNase-A were added to each tube and incubated overnight at 65 °C. Total DNA for fruits and feces was extracted using the Qiagen® DNeasy Plant Mini Kit according to the manufacturer’s instructions.

PCR amplicons of the ITS2 nrDNA region using fungal-specific primers fITS7 and ITS4 (Ihrmark *et al.* 2012) were tagged and multiplexed for paired-end sequencing (250 bp) on the Illumina MiSeq platform at MRDNA (http://mrdnalab.com, Shallowater, TX, U.S.A.). Three PCR stochastic replicates were done and then pooled. Pooled samples were purified using calibrated AMPure XP beads (Beckman Coulter Life Sciences, Indianapolis, IN, U.S.A.). PCR from a pure fungal culture of a *Trichoderma koningiopsis* was used in quality control in the downstream bioinformatics. Sequencing depth was at approximately 160K reads per sample.

### Bioinformatics and fungal species identification

We used MOTHUR v 1.36.1 (Schloss *et al.* 2009) to assemble paired-ends. FastQ files were then imported into Geneious v10.0.7 (Biomatters Ltd., Auckland, New Zealand) to trim primers from contigs and low-quality sequences or regions (error probability limit = 0.05). All sequences with length equal to zero were also deleted in Geneious. USEARCH v10 (Edgar 2010) was used to trim sequences to equal length of 200 bp, to delete sequences with a maximum number of expected errors of 0.5, to dereplicate sequences, and to delete singletons. Sequences were clustered into 99%-similarity Operational Taxonomic Units (OTUs) (Lim *et al.* 2010; Gazis *et al.* 2011; Ikeda *et al.* 2014). After clustering, OTUs were subjected to similarity searches in the UNITE curated and quality-checked database (Kõljalg *et al.* 2013). Sequences were filtered for chimeras using the latest ITS chimera dataset (Nilsson *et al.* 2015) and implemented in UCHIME (Edgar *et al.* 2011). UNITE assigns *species hypothesis* (SH) to each OTU. OTUs were assigned to a genus, family, order, class or phylum using similarities between 95-98%, 90-95%, 85-90%, 80-85%, and 77-80%, respectively. OTUs with less than 77% were labeled as *incertae sedis,* and anything below 70% was deleted from further analyses. UNITE similarity searches result in 10 species hypotheses for each OTU. Taxonomic accuracy was checked manually for each of the 10 SH for each OTU because in some cases the best match may be incorrectly identified and named in the database, and also to standardize fungal names and avoid synonyms. Then, each name was verified for correctness either in Index Fungorum or Mycobank. Raw data, fastQ files, are available upon request.

A total of 5,915,682 reads were obtained after quality filtering and trimming with an average read length of 322bp (±SD = 21bp). Libraries for each sample varied between ca. 1400 and 230,000 reads. Due to the large variation in reads per sample, we normalized the data in R by subsampling to 1408, the lowest number of sequences amongst all samples. These data were then used for community analyses. In other cases, non-normalized data was used for descriptive purposes and is indicated.

### Fungal diversity and community analyses

To characterize fungal communities in bat feces and fruits, we first calculated species richness (S), diversity (Shannon, H), and evenness (H/lnS) based on the normalized abundance data in Qeco (Quantitative Ecology Software, (Di Rienzo *et al.* 2010). These values were estimated for each fruit (n = 9) and bat feces (n = 13) sampled, and were later compared with a Mann-Whitney U test in InfoStat version 2017p (Di Rienzo *et al.* 2017). We also used rarified abundance data of OTUs to calculate the expected species richness and sample coverage, based on Hill numbers (q=0), using a combination of rarefaction and extrapolation; we extrapolated to twice the reference sample size (Chao *et al.* 2014). The analyses were further repeated with a matrix of presence-absence data for confirmation of results.

Fungal communities in feces and fruits were compared based on rarified abundance and presence-absence data. We used an Analysis of Similarities (ANOSIM; Clarke 1993) in Qeco, implemented from the package vegan (Oksanen *et al.* 2013). ANOSIM tests the null-hypothesis that the similarity between communities (i.e., bats and fruits) is greater than within communities. The test statistic R may take values between −1, and 1; negative values suggest there are greater similarities between communities, whereas positive values suggest greater similarity within communities (Clarke 1993). To determine similarity, we used the Bray-Curtis distance measure on standardized data, as some species were very abundant and others were very rare. For presence-absence data, we used the Jaccard distance measure. We used 1000 permutation cycles for calculating a p-value. Additionally, we identified indicator value species, IVS (De Cáceres *et al.* 2010), or those that may reflect the biotic or abiotic state of the environment and predict the diversity of other species within a community (McGeoch 1998; Niemi & McDonald 2004). The species (or OTUs) were selected as indicators of either the bat or fruit fungal assemblages if they showed a high indicator value and significant p-values for the corresponding community.

### Estimation of abundance of fungal taxa according to their ecological roles

We explored if there was a trend in the abundance of OTUs with specific ecological roles in fruits and feces. A putative ecological or functional role was assigned by parsing fungal community datasets by ecological guild using FUNGuild annotation tool (Nguyen *et al.* 2016) and then manually checking and confirming the assignments (following the approach in Gazis & Chaverri 2015). OTUs were assigned to one of the following putative roles: (1) animal gut endosymbiont, (2) insect endosymbiont or entomosymbiont, (4) mycotroph, (5) plant pathogen, (6) insecticolous or entomopathogen, and (7) plant saprobe.

With the data on ecological roles and abundance for taxa, we then constructed a heat map using the package gplots in R. For this map, we only included the most abundant taxa, i.e., those for which more than 100 hits were recorded. First, we normalized abundance data by dividing the number of hits for each taxon over the total abundance for a specific sample (i.e., bat or fruit). The heatmap was created by plotting the normalized abundance of each taxon for each sample, and the similarity of fungal communities among samples, shown in a dendrogram, was estimated based on Euclidean distance.

## RESULTS

Due to the rarity of the bats, the difficulty to find several fruiting trees within our study sites, and the low probability of capturing bats near those fruiting trees, our study is based on results of several fruits collected from two trees and thirteen bats, which is still a sufficiently large sample size to answer our study questions. By capturing bats near a fruiting tree, we were able to increase the probability that the fecal samples came from fruits consumed from those two trees, based on previous observations that bats may exclusively feed on a single tree throughout the night and even during several consecutive nights (Villalobos-Chaves *et al.* 2017).

The total number of fungal OTUs identified from metabarcoding from 9 fruits and 13 bats was 188 and 388, respectively (supplementary Tables S1–S4). Fungal communities in bats exhibited higher species richness (mean = 48.77, ±SD 23.45), greater evenness (0.63 ± 0.20), and higher diversity (2.43 ± 0.95) than those in fruits (richness: 42.33 ± 15.71, evenness: 0.47 ± 0.21, diversity: 1.74 ± 0.88), yet the difference between them was not significant (p-values for the Mann-Whitney U test > 0.07). Our sample size was not sufficient to detect the expected number of species for fruits and feces. Species accumulation curves (Fig. 1a) show that an asymptote in species richness (based on Hill numbers; q=0) was not achieved for both fruit and bat feces assemblages when we extrapolate to at least double the current sample size. In addition, estimates of sample coverage (Fig. 1b) show that our current sampling effort represents only 0.46 and 0.35 sample coverage for fruits and bat feces, respectively. If we extrapolate to a sample of 18 for fruits and 26 for bats (double our current sample size), sample coverage only slightly increases to 0.62 for fruits, and 0.50 for bat feces. Results were similar when comparing communities using the presence-absence data (Table S5, Fig. S1).

**Figure 1.**
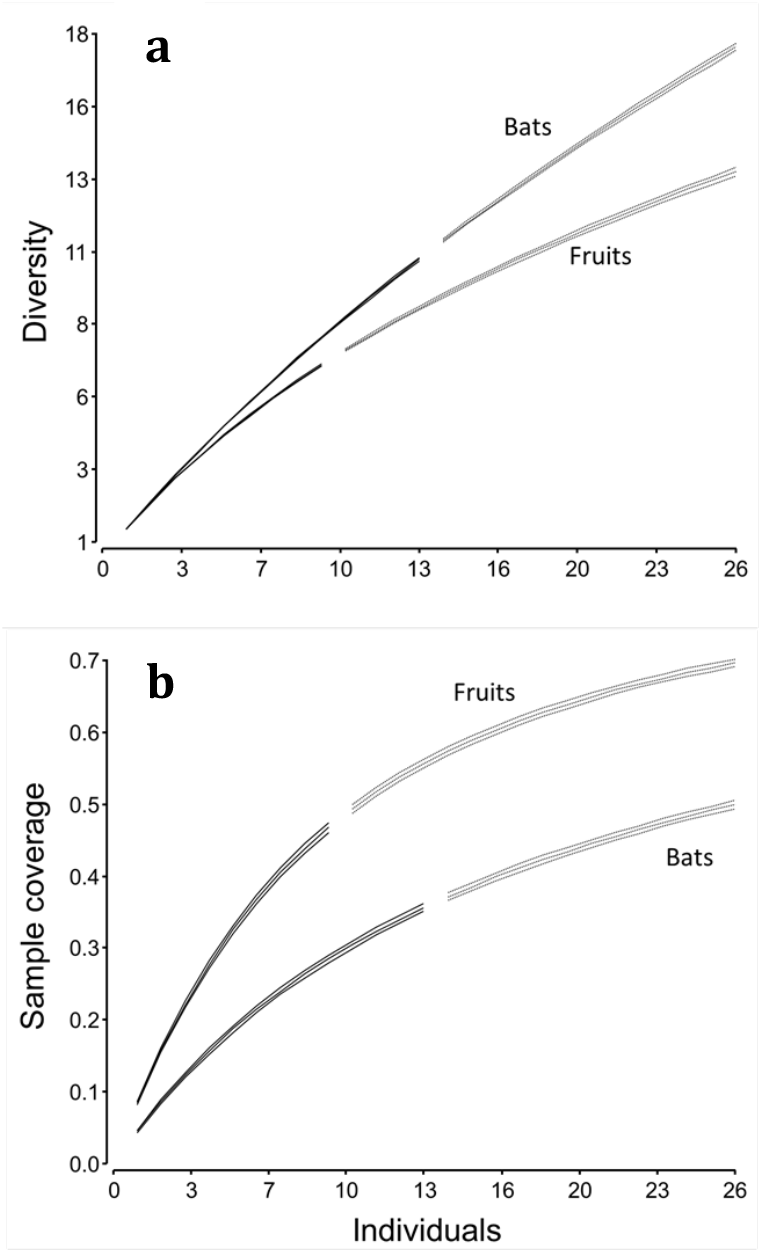
Sample-size-based rarefaction (first set of solid lines) and extrapolation (second set of dotted lines) of fungal diversity (based on Hill numbers; q=0) in bat (*Ectophylla alba*) feces and fruits (*Ficus colubrinae*). External lines represent the 95% confidence intervals obtained from 50 bootstrap replications. a) Diversity; b) Sample coverage.

ANOSIM, using the rarified abundance data, shows that fungal communities in fruits and feces were significantly different (R=0.21, P<0.001); results for presence-absence data indicate borderline significant differences between the two communities (R=0.12, P=0.06). The fungal species with greater indicator values (>0.80) for fruits included an unidentified family within the order Pleosporales, two species within the genus *Phyllosticta*, *Colletotrichtum citricola* and *Edenia gomespompae* (Table 1). For bat feces, only two fungal species had indicator values >0.80, namely *Cladosporium cladosporioides* and *Alternaria alternata*. Interestingly, many of the indicator species in bats with values >0.70 belong to the genus *Aspergillus*. Results of IVS were similar for presence-absence data (Table S6), with the addition of three taxa as indicators of fruit communities (*Powellomyces* sp., *Phlebia tremellosa*, and *Diaporthe* sp.) and exclusion of others (*Edenia gomezpompae* in fruits, *Alternaria alternata* in bats).

**Table 1.**
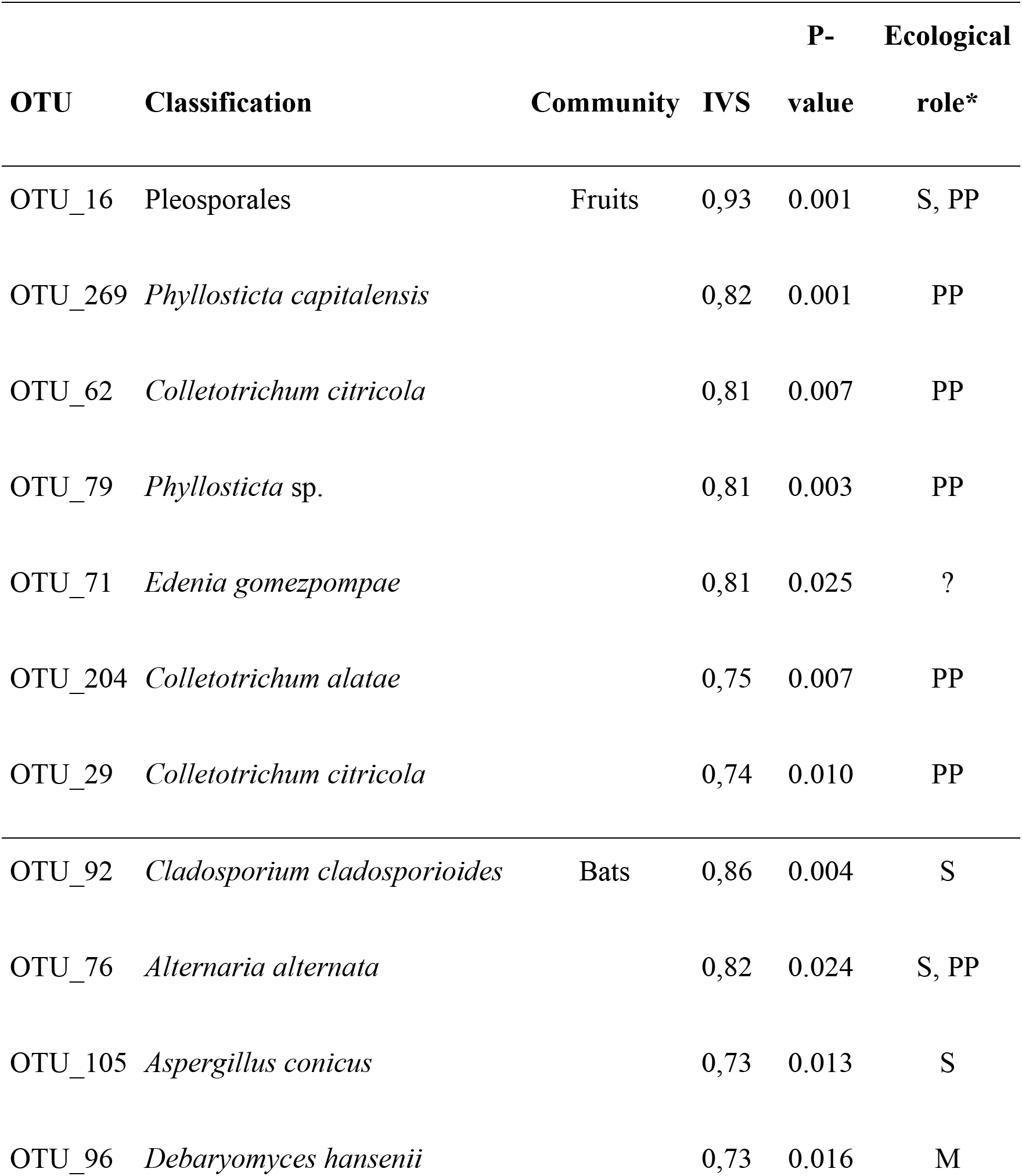
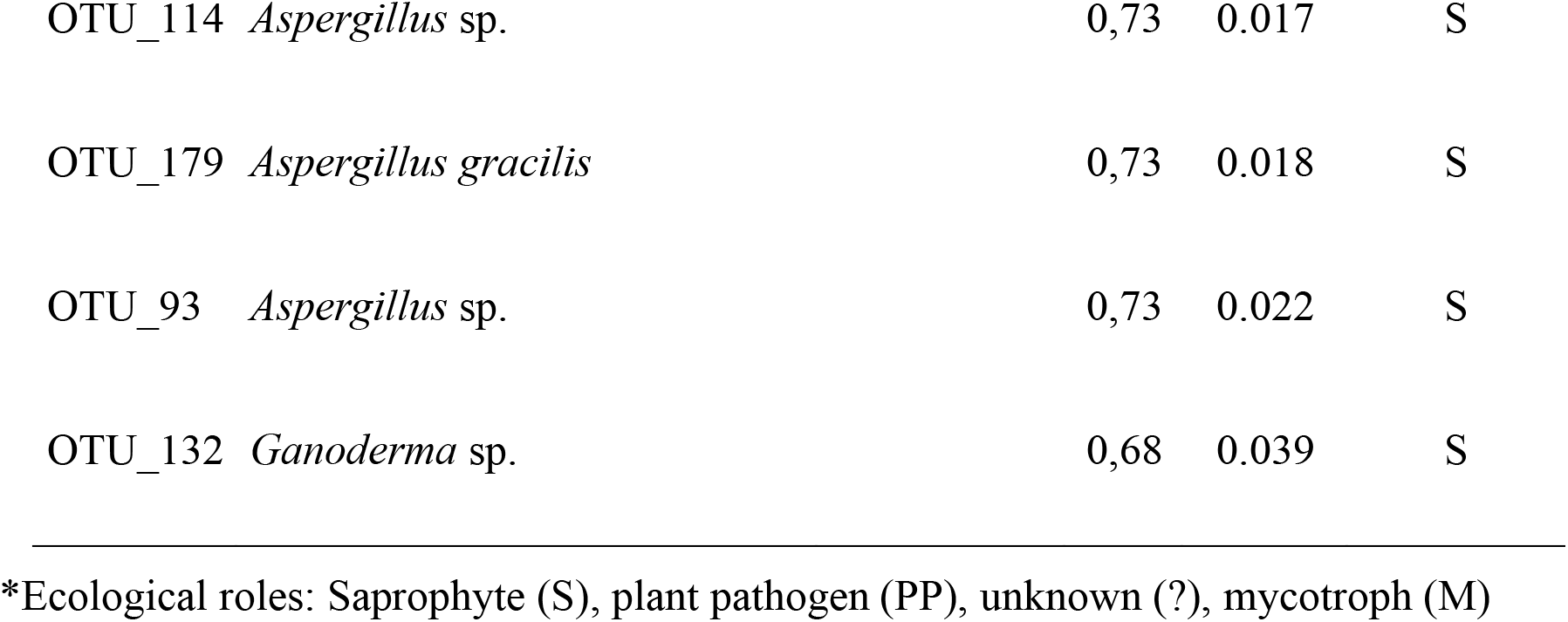
Indicator value species (IVS) and associated p-values for the endophytic fungal assemblages in the two communities studied, fruits (*Ficus colubrinae*) and feces in bats (*Ectophylla alba*). We provide species names wherever possible, or alternatively the closest classification possible for a given OTU. Ecological roles are included.

Animal commensal/gut endosymbionts were more abundant in feces, and plant pathogens dominated the fungal community in fruits (Fig. 2). Plant saprotrophs were abundant in both fruits and feces. Our results also show that some of the most common OTUs in fruits were almost absent or rare in feces, and vice versa (supplementary material Tables S1–S4, S7, and Figs. S2, S3). With the normalized data, the most abundant fungal orders (>100 hits) in fruits were Pleosporales, Glomerellales, and Hypocreales (Fig. 3). The most abundant orders in bat feces (>100 hits) were Hypocreales, Wallemiales, Malasseziales, Agaricales and Cystofilobasidiales (Fig. 3) using normalized data. The non-normalized data showed a similar trend for both fruits and bat feces (Fig. S4). More specifically, *Colletotrichum* spp., *Corynespora cassiicola*, *Curvularia lunata*, *Diaporthe* spp., *Edenia gomezpompae, Fusarium* cf. *decemcellulare, Phyllosticta* spp., and *Wickerhamomyces queroliae,* which were very abundant in fruits, had a >90% reduction in abundance in feces (supplementary Tables S1–S4, S7). On the other hand, *Cladosporium* cf. *cladosporioides, Cystofilobasidium* spp., *Diplodia seriata, Exobasidium* spp., *Fusarium* cf. *proliferatum, Malassezia* spp., *Phlebia tremellosa, Powellomyces* sp., *Sporobolomyces* spp., and *Wallemia* spp., among others, were 70-300 times more abundant in feces than in fruits (supplementary Tables S1–S4, S7).

**Figure 2.**
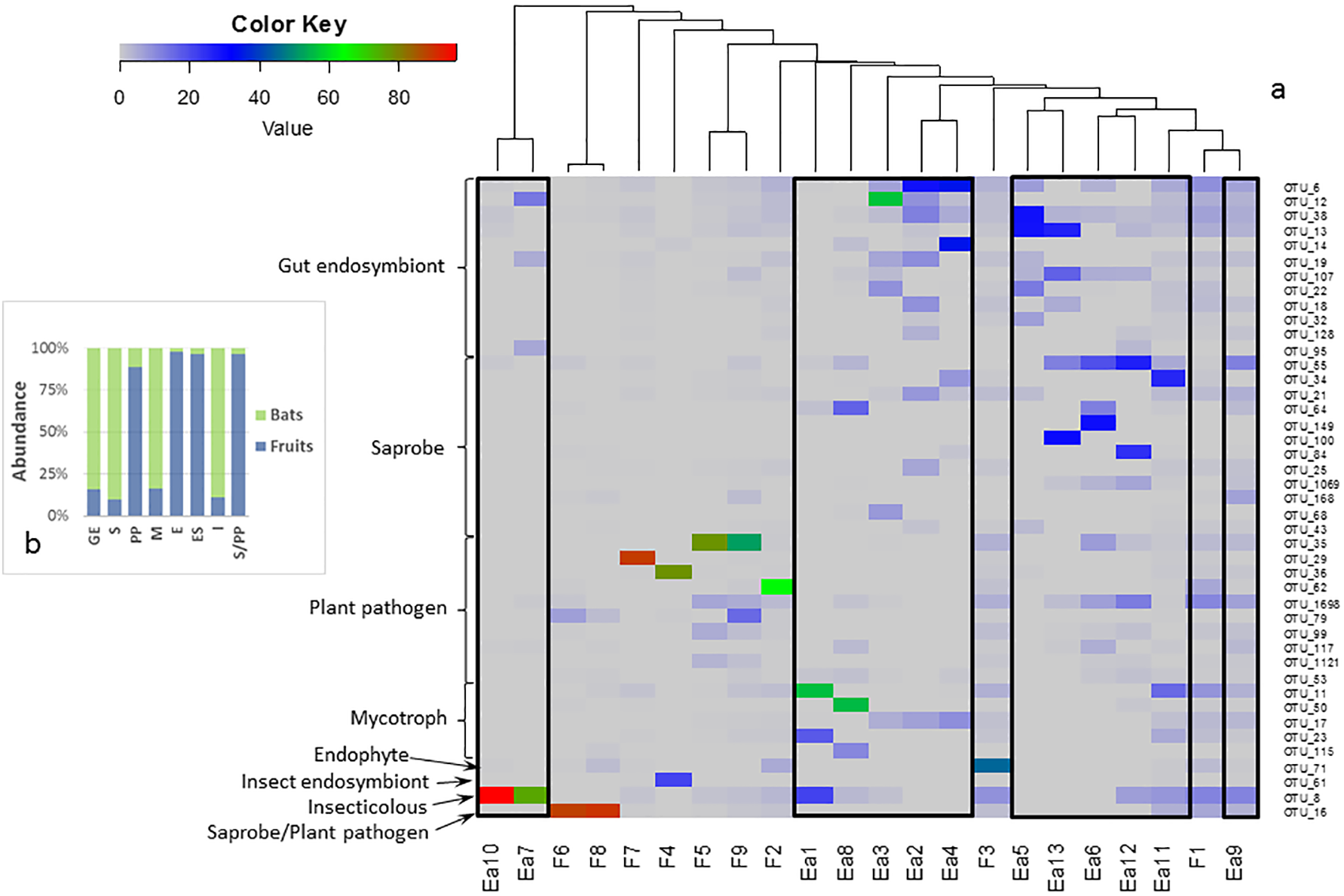
a) Heatmap that shows the abundance of fungal taxa for which more than 100 hits/reads were recorded (right labels) in bats and fruits (bottom labels; bats: Ea, fruits: F). Fungal taxa are separated by ecological roles (left labels), and samples are arranged according to the similarity (top dendrogram) in their fungal communities. Boxes encompass samples from bats. b) Graph that summarizes the differences in total abundance of fungal taxa between fruits and bats grouped by ecological roles (labels follow the same order as a, top to bottom). Normalized data was used for this heatmap.

**Figure 3.**
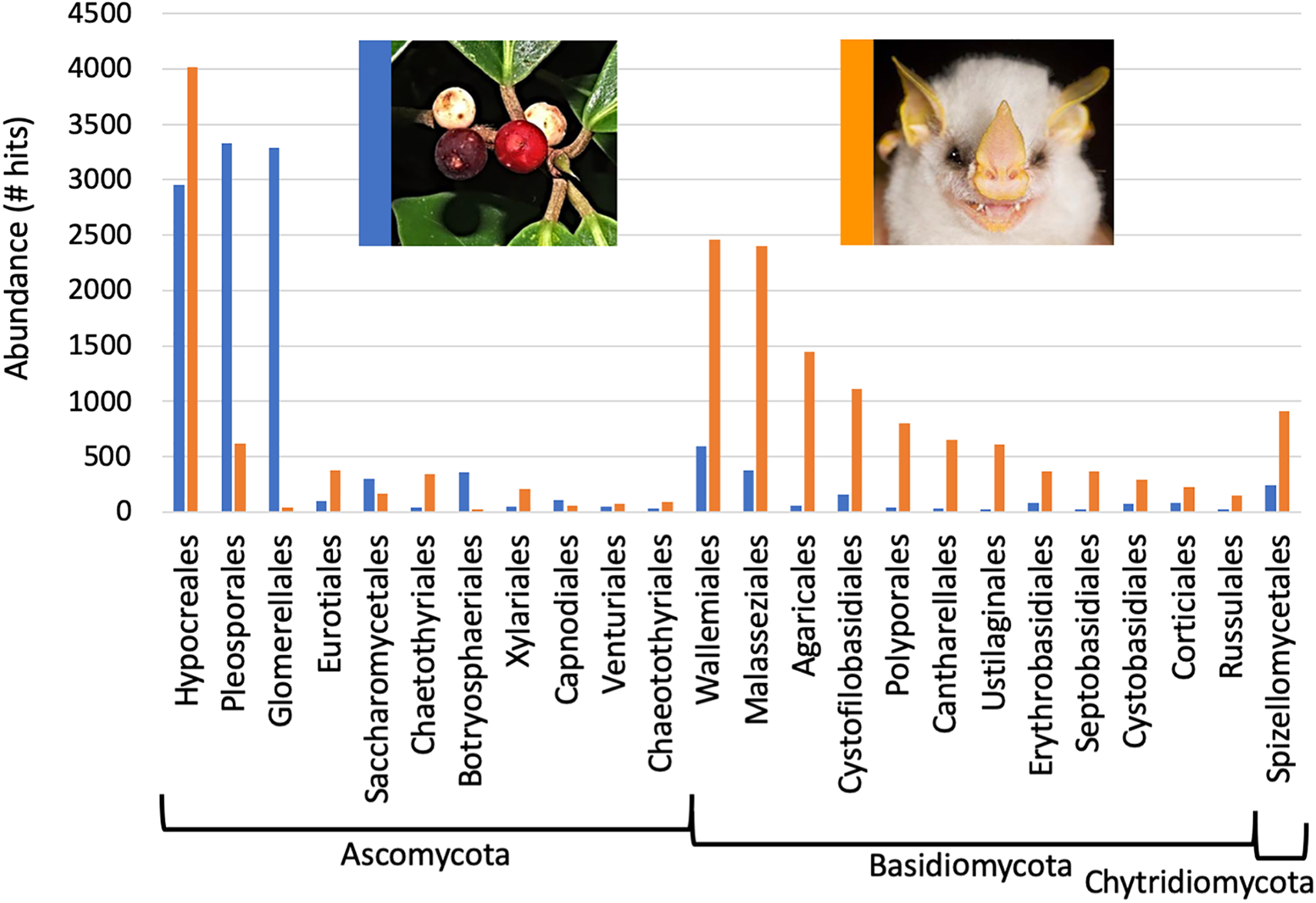
Relative abundance of the 24 most abundant fungal orders in fruits of *Ficus colubrinae* (blue bars) and feces of *Ectophylla alba* (orange bars). The vertical axis represents the number of hits/reads with the normalized data. Classification of each OTU is detailed in supplementary Table S10.

## DISCUSSION

The results of our study show a diverse epi- and endosymbiotic fungal community in fruits (188 observed spp. and >400 expected) and bat feces (388 obs. and >900 exp.). Similarly speciose communities have been observed in other plant tissues, such as leaves, roots, stems, and sapwood (Arnold *et al.* 2001; Vega *et al.* 2010; Gazis & Chaverri 2015), and several studies confirm that many of the fungal species identified thus far provide benefits to their plant hosts (Arnold *et al.* 2003; Gazis & Chaverri 2015). Interestingly, our study suggests that while fungal communities are quite diverse in fruits, with several potentially pathogenic as well as beneficial taxa, by passing through the bat’s gut the former tend to become rare, while the latter more abundant. This change in the mycobiome from fruits to feces is supported by our results of community analyses and by substantial changes in abundance of certain taxa (supplementary Table S7). Most striking was the reduction of >70% in abundance of taxa known to be plant pathogenic (or hemibiotrophic) after passage through the gut. In contrast, species known to have antifungal and antibacterial properties, or to provide protection against UV-B light and even nutrition to bats (e.g., carotenoids, lipids, coenzyme Q10), significantly increased (supplementary Tables S8 and S9). These findings suggest that the role of frugivores in plant-animal mutualistic networks may extend beyond seed dispersal: they may also promote the dispersal of potentially beneficial epi- and endosymbionts while hindering those that can cause disease.

Our characterization of the mycobiomes indicates the presence of fungal DNA (i.e., ITS nrDNA) in fruits and then in bat feces, yet we can only confidently conclude that there is presence of fungal DNA, but not that the fungi in feces are still viable. Therefore, we cannot be certain that DNA from many beneficial fungi present in feces has the capacity to germinate. Even though we did not culture fungi from fruits and feces, results from other studies indicate that several fungal taxa can grow after passing through the digestive tract of vertebrates (Johnson 1996; Bertolino *et al.* 2004). Additional RNA, metatranscriptomic or proteomic analyses may reveal more about the viable mycobiome, in addition to functions and interactions between microbial species (Sørensen *et al.* 2009; Kirschner *et al.* 2015; Spain *et al.* 2015; Zhao *et al.* 2015).

Comparing the mycobiomes of fruits and bat feces provides seminal results on how frugivores may be affecting plant tissues that are exposed to microbial communities within feces, how surviving fungal taxa may affect frugivores, and how fungi may affect the interaction between fruits and frugivores. Dispersed seeds may benefit from frugivores by a reduction in the number of potentially pathogenic taxa and an increase in the numbers of protective fungi to which they are exposed to when deposited in the soil as a result of passing through an animal’s gut. These interactions could increase seed survival, which is often very low primarily due to attacks by pathogens (Dostál 2010). We can infer that if *F. colubrinae* fruits are not consumed by frugivores and they just fall from the tree, potential plant pathogens contained in fruits will remain near the parent tree and cause disease in seedlings. These results support the Janzen-Connell hypothesis (Connell 1971; Janzen 1971) and the Theory of Pest Pressure (Gillett 1962), which suggest that specialized natural enemies (i.e., plant pathogens) decrease survival of seedlings that are in high densities beneath the parent tree, thus giving locally rare species an advantage. An alternative scenario is that if fruit-eating bats defecate under or near *F. colubrinae* trees, or under or near their roosts (leaves of *Heliconia* spp.) (Morrison 1980), passage through the digestive tract still results in decreased abundance of potential plant pathogens, which would reduce the amount of plant disease. Therefore, consumption of fruits by animals (i.e., bats) not only benefits the plant by dispersing its seeds, but also by reducing the amount of pathogen inoculum and thus escaping disease pressure (Gilbert 2005). On the other hand, seedlings may incorporate a beneficial fungal community into their own tissues since many potentially beneficial taxa (e.g., anti-herbivory and antimicrobial) survive and increase in abundance after passage through the digestive tract of *E. alba* (supplementary Tables S7 and S8).

While we still lack sufficient information to predict how certain fungal taxa found in the tissues of fruits may be affecting fruit-eating animals, some of the fungi we recorded in feces that were also present in fruits are considered common animal endosymbionts associated to healthy guts (Kunčič *et al.* 2010; Amend 2014), such as Malasseziales and Wallemiales. Finally, fungi may also affect the interaction between plants and frugivores. It is known that fungi can promote the synthesis of secondary metabolites (Venugopalan & Srivastava 2015), which can affect fruit palatability (Whitehead & Poveda 2011). By extension, we hypothesize that endophytic fungal communities can change the chemical composition of fruits and hence preferences in frugivores. While unquestionably relevant for understanding mutualistic networks, this topic remains completely unexplored.

This poorly known interaction among fungal endophytes, fruits, and frugivores suggests there is a critical component of plant-animal mutualistic networks that urgently requires further scrutiny. This additional link complicates our understanding of mutualistic network dynamics and perhaps many of the models developed thus far may not fully account for the effect of the loss of a particular species on those networks. For example, we often regard frugivores as somewhat equally responsible for promoting seed dispersal, yet differing foraging styles and physiological conditions among fruit-eating species can potentially affect dispersal distance and viability of propagules, fungal or otherwise (Colgan & Claridge 2002; Charalambidou *et al.* 2003). By affecting fruit palatability and overall plant fitness, the mycobiome may also be highly responsible for the success, or failure, of certain plant species, which ultimately modifies the composition of many ecological networks and interactions therein.

The study of the mycobiome has experienced a major increase in the last years, predominantly with the advent of genetic tools, and evidence is accumulating on the many roles that fungi play in natural ecosystems. Many of the studies conducted so far have shown a diverse fungal community in plants, yet surprisingly, very little is known about mycosymbionts in fruits (but see Vega *et al.* 2010; Tiscornia *et al.* 2012; Paul *et al.* 2014; Taylor *et al.* 2014), despite the obvious role of fruits for plant fitness, and no studies to date have assessed how vertebrate consumption can affect fungal endophytes. As a consequence, the role of endophytic fungi in mutualistic networks has been, until now, largely ignored. Our study shows that fungal endophytes are ubiquitous within fruits, and as such may be important components of plant-animal networks. Our study also suggests that frugivores may play yet another essential role within these networks by hampering pathogenic, and enhancing beneficial, fungi dispersed within their feces. Their ubiquity in plant tissues, and the potential role that plant-eating organisms can play in dispersing the fungal propagules, suggest that further studies into mutualistic interactions should greatly focus on endosymbionts.

## ACKNOWLEDGEMENTS

We deeply acknowledge J.P. Barrantes, C. Castillo-Salazar, and E. Hellman for their help in laboratory and fieldwork. Joxerra Aihartza, Inazio Garin, Lide Jiménes, Edwin Paniagua, and Emmanuel Rojas also provided valuable field assistance. Aline Vaz also provided assistance in creating a heatmap and József Geml with rarefaction. Finally, we thank the staff at Tirimbina Biological Reserve and La Selva Biological Station for their help with logistics and accommodation, and Lourdes Vargas from SINAC for her help with research permits. This research was supported by a grant from Conservation, Food and Health Foundation.

